# FIRST EVIDENCE OF OBJECT PLAY IN WILD GELADAS: FUNCTIONAL IMPLICATIONS FOR LATER UTILITY AND RE-ELABORATED OBJECT USE IN ADULTHOOD

**DOI:** 10.64898/2026.03.23.713729

**Authors:** Giada Cordoni, Maria Chiara Porfiri, Yibelu Yitayih Hailie, Anna Benori, Sebastiano Bergamo, Ejigu Dessalegn Berhane, Bezawork Afework Bogale, Ivan Norscia

**Affiliations:** Department of Life Sciences and Systems Biology, University of Torino (Turin, Italy); Department of Biology - College of Science, Bahir Dar University (Bahir Dar, Ethiopia); Department of Biological Science - College of Natural and Computational Science, Addis Ababa University (Addis Ababa, Ethiopia)

**Keywords:** social object play, solitary object play, behavioral ecotypes, *Theropithecus gelada*, *Papio anubis*

## Abstract

Object play - seemingly non-functional interactions with objects – can promote the development of foraging skills, tool use, and behavioral innovation. Among Catarrhine monkeys, it was described in macaques and baboons. Wild geladas, although closely related to baboons, have been described as lacking object play (observed only in captivity) linked to their specialized grazing ecology. Here, we provide the first evidence of both social and solitary object play in a wild gelada population (N_OMUs_=13) at Debre Libanos (Ethiopia) and compare it with object play in sympatric olive baboons (N_individuals_=42). Notably, immature geladas engaged in object play both socially and solitarily, but the latter case was most frequent also with novel objects introduced by researchers. Solitary object play occurred at levels comparable to those of baboons, challenging previous reports of limited object interest in geladas. This finding aligns with the occurrence of object play in phylogenetically related species and with the retention in wild geladas of arboreal behavior and fruit consumption and hand morphology enhancing fine manipulation. Hence, object play in geladas under certain environmental conditions may reflect a biologically rooted capacity and underscores the importance of ecological variability, as distinct ‘*behavioral ecotypes*’ can emerge across different populations of the same species.

---

Object manipulation in animals can occur via play and exploration [1–2]. For a long time, play and exploration were considered two similar phenomena, differing only in their intensity [3]. Recently, it has been proposed that exploration provides information about an object’s affordances (i.e. “*what can this object do*?”), whereas play allows individuals to discover the *creative* use of an object (i.e. “*what can I do with this object*?”) [1]. In this sense, object play involves elements of exploration, but not all exploratory behaviors necessarily constitute play [2].

Children play with objects in a modified form compared to the way those same objects, or more functional versions of them, are used in adult life [4]. Object play (or play-like behavior) is not limited to humans, but is also found in other animals, ranging from invertebrates to vertebrate mammals (e.g. octopuses [5], cichlid fish [6], lizards [7], crocodilians [8], corvids and parrots [9–10], kittens [11], and dolphins [12]). Indeed, object play is evolutionarily relevant, as it can modify ontogenetic contexts and shape selection pressures, in line with the niche-construction theory [4]. During object play, which goes beyond mere exploration [1,2], individuals physically engage with objects by picking them up and combining their use with other objects and/or elements of the environment [13–15]. Although object play appears to provide no immediate benefit [1–2,14–15], it helps individuals navigate social relationships over resources - especially when social - and acquire knowledge of the action-relevant properties of objects while enhancing manipulative abilities (such as foraging skills) - especially when solitary [1–2,15–16]. Importantly, object play is also a precursor to re-elaborated tool use later in life and can generate behavioral innovation by fostering novel object use [1,17–21].

Complex object play is especially found in primates (e.g. great apes, [17,22–24]). For example, among great apes, wild immature chimpanzees perform many atypical uses of objects during play - including a use that is different from adult uses (e.g., dipping stick into water to attempt drinking) - that meet the criteria for innovations because they are individually generated, spontaneous, and novel forms of behavior [21]. Among Catarrhine monkeys, macaques engage in stone handling which represents a flexible and complex form of object play [17,25–27]. Notably, some playful stone-handling patterns may have been co-opted into the use of stones as tools in other contexts, such as aggression or feeding [17,25]. Furthermore, both captive and wild baboons (genus *Papio*) engage in object play and show a marked interest in novel objects, which may be used as tools for food processing and extraction during adulthood [28–29].

Unexpectedly, geladas (*Theropithecus gelada*) - the Catarrhine monkey species phylogenetically closest to the genus *Papio* [30–31] - do not interact with novel objects in the same way as baboons [29]. Moreover, geladas have been reported to play with objects in captivity (stone handling) but not in the wild [32]. This difference has been linked to the fact that baboons live in a wide range of habitats (e.g. grasslands, savannas, mountains, rainforests) and consume diverse types of food [33–36], whereas geladas are mostly herbivorous and largely restricted to the highlands of Ethiopia [37–38]. However, gelada ecology is more complex than it may appear. Geladas inhabit diverse habitats, including both grasslands and rocky cliffs [37–38], the latter preventing full observation, which limits our knowledge of their eco-ethology. They also retain some arboreal habits and frugivory [39–40], traits deeply rooted in primate biology since the Eocene [41]. They possess short digits, a trait shared with baboons, which enhances thumb-hand integration and thereby supports complex manipulative abilities [42]. Moreover, gelada hand anatomy is the closest to that of humans among Catarrhines, and during feeding they use pad-to-pad precision grips, in-hand movements, and compound grips, all among the most advanced manual skills in primates [43].

Based on these elements, it is counterintuitive that object play in geladas - despite the species possessing the biological hardware and all the prerequisites to express it - would be completely absent in the wild. We hypothesized that, under certain environmental conditions, object play in geladas may occur and may be adaptive, as in other phylogenetically related species, by facilitating the later emergence of re-elaborated object use in adulthood. To test this hypothesis, we describe and analyze social and solitary object play for the first time in wild geladas, comparing it with object play in sympatric baboons and with object use in adults.

## METHODS

### The study area and groups

The study was carried out on wild geladas (*Theropithecus gelada*) and olive baboons (*Papio anubis*) inhabiting the Fiche zone, Debre Libanos Woreda, Debre Tsige town, Addis Alam village, Set Deber district (9.42471° N; 38.51257° E). The area was surrounded by croplands, monastery, and village (Set Deber) on three sides and by rocky cliffs on the remaining side. The area frequented by baboons and geladas was anthropized and characterized by the presence of discarded items left by people.

#### Geladas

For the purpose of this study, 13 One-Male Units (OMUs; N_individuals_=175) were considered. The mean number ±SE of adult and immature individuals (i.e. infants, juveniles, and subadults) per OMU was 8.4±1.5 and 4.5±1.0, respectively. OMUs occupied the Debre Libanos plateau and their pasture area consisted of the grassland portions without human settlements, where livestock (horses, goats, sheep, donkeys, and cows) grazed during the day under the supervision of shepherds. Gelada groups freely moved up and down the cliffs from the plateau.

#### Olive baboons

One multi-male/multi-female troop comprising a total of 42 individuals (17 immature individuals and 25 adults) was observed. The troop frequented an area overlapping with the highlands frequented by geladas but also included a forested area along the river, around and within the cemetery area.

Geladas and baboons were individually identified by focusing on distinctive morphological and behavioral features [37,44–45]. The identification of adult males in geladas allowed OMU identification.

### Data collection and operational definition

Live and video data were collected from February to June 2025 on both geladas and olive baboons. Observations were carried out 5 days per week, from approximately 08:00 am to 02:00 pm for security reasons. Video recordings were obtained using full HD cameras (Lumix Panasonic DC-FZ82, zoom 60x, 30fps; handy-cam Panasonic HC-V180, zoom 90x, fps). In total, 108 hours of video were collected for geladas and 54 hours for baboons.

We employed scan animal sampling (live data; [46]) at 15-min intervals across all OMUs and the baboon troop to record feeding, moving, resting, grooming, aggression, and solitary and social play interactions. In addition, we used all-occurrences sampling to document all solitary and social play events in both species. For each event, we recorded the identity of the player(s) and the presence and type of object used during play.

A play session – distinguishable from other behavioral contexts by the presence of play signals (e.g. play face and full play face), the repetition and exaggeration of the patterns performed, and the absence of threatening and/or fear signals (e.g., scream face, avoidance) [1–2,14–15] - was defined as social if it involved two or more individuals; otherwise, it was defined as solitary.

A play session began when an individual performed a playful pattern (see Table 1) alone or directed one toward a conspecific and ended when play activity stopped for more than 10 s or when another individual joined or replaced one of the players (thereby initiating a new session) [47].

**Table 1.**
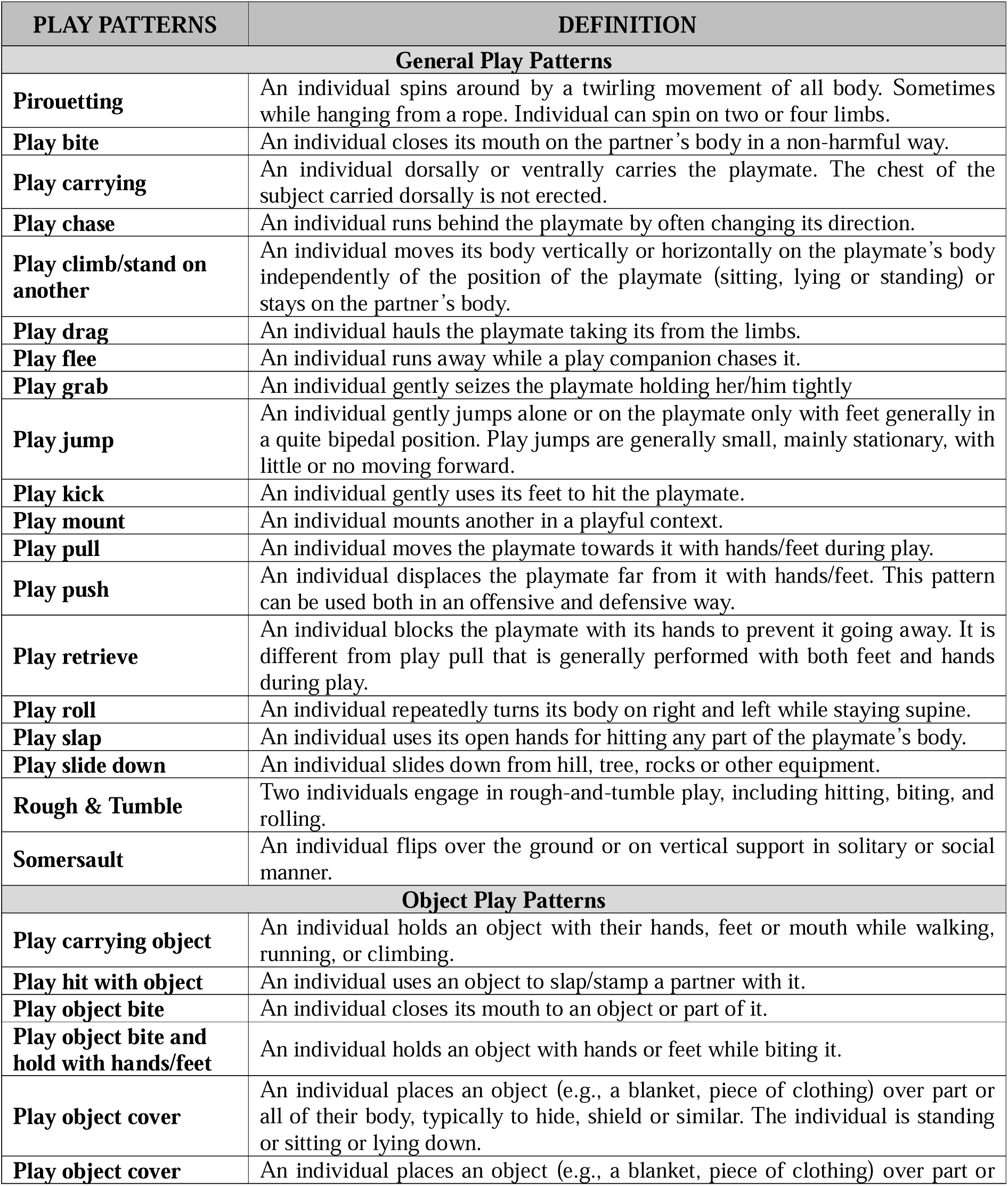
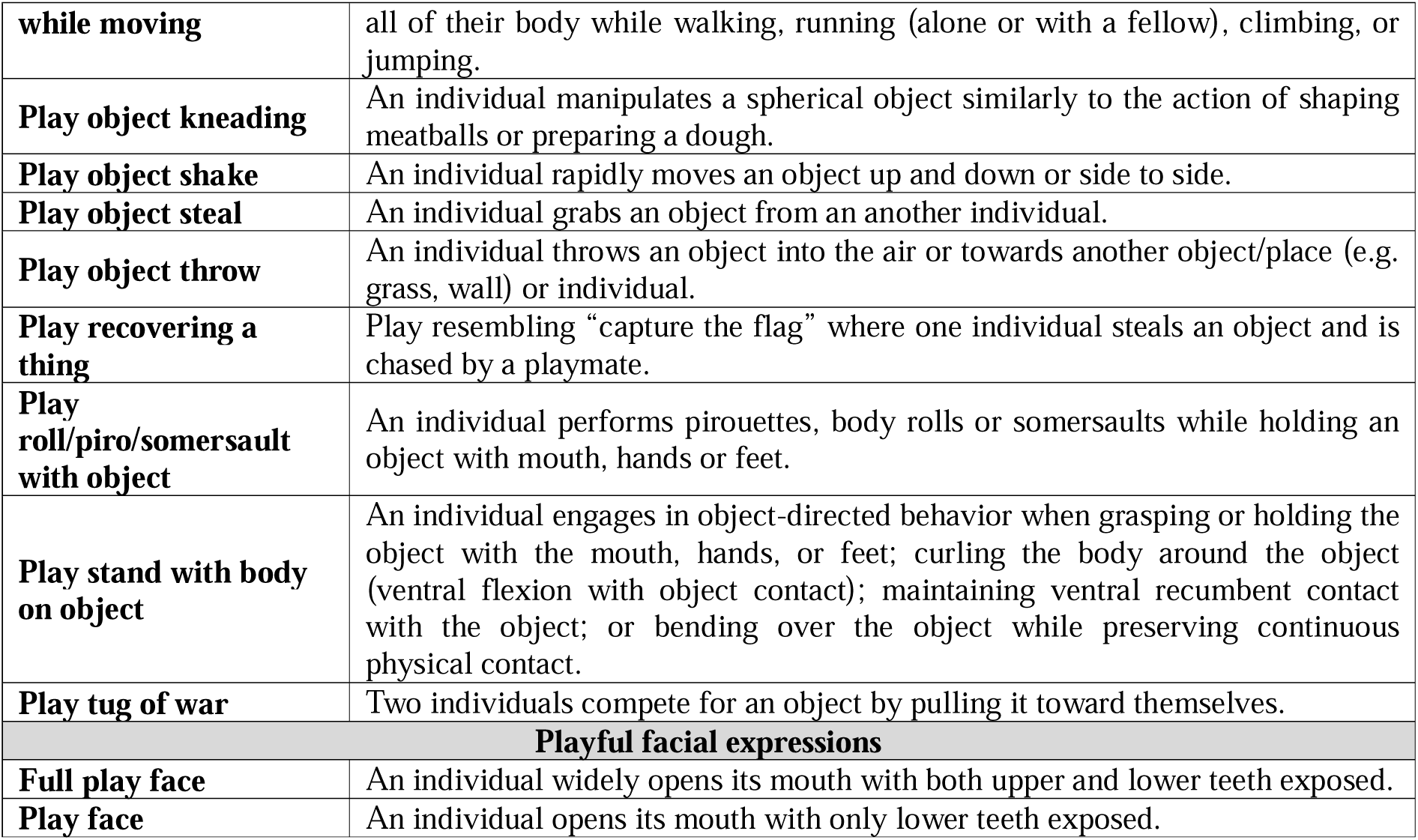
Play behavioral patterns of both geladas and baboons considered in this study.

The researchers provided geladas with novel objects (which are not present in the environment) on specific days alternated with days in which no objects were provided; this procedure was adopted to avoid habituation effects and to also observe naturally occurring use of objects. The researchers distributed clusters of colored plastic balls (diameter 7 cm each) and colored ropes (length 1 meter each) across the plateau. Balls and ropes are valid shapes in adaptive and ecological terms as spherical/subspherical fruits and vines are common in primate environments. The temporary provision of objects was intended to increase the likelihood of observing object play, given that the presence of novel manipulable items can enhance play behavior [19–21].

### Statistical elaboration

For each individual, we calculated a Social Object Play Index (SocOPI) and a Solitary Object Play Index (SolOPI) as the number of social or solitary play sessions involving an object divided by the total number of social or solitary play sessions, respectively. For geladas, we excluded the sessions where the objects had been provided by the researchers.

Due to the non-normality of data (Kolmogorov-Smirnov test, 0.197<Z<0.500, p<0.001), we compared SocOPI and SolOPI between geladas and baboons using the non-parametric Mann–Whitney test for two independent samples. We then used the Wilcoxon signed-rank test for paired samples to compare: i) SocOPI and SolOPI within each species; ii) and to compare the frequency of social and solitary object play in geladas only considering the cases where objects were provided by researchers. Montecarlo procedure (10,000 permutations) was applied to account for non-identified individuals.

## RESULTS

During the observation period, in geladas, we recorded 133 social play sessions, 8.3% of which involved an object, and 31 solitary play sessions, 48.4% of which involved an object. Playful sessions performed when objects were provided by the researchers were excluded from this count. Only immature individuals engaged in social and solitary object play.

In olive baboons, we recorded 1.413 social play sessions, 7.1% of which involved an object, and 222 solitary play sessions, 43.7% of which involved an object. Among the object play sessions, adult baboons were involved in 10 social and 14 solitary sessions.

During object play sessions (both solitary and social), geladas used sticks/branches (Fig. 1a; video S1), plastic bottles, parts of clothes, waste, and shoes (Fig. 2a; Video S2). We also observed 17 aggression (8 of which involved subadult individuals) during which opponents employed objects like stones, sods of earth, wooden stakes, parts of clothes, waste, and branches (Fig. 1b; Video S1). Notably, when researchers provided objects not typically present in the geladas’ environment, immature individuals also played with plastic balls and ropes (Fig. 2b; Video S2). Similarly, baboons used plastic bags and bottles, clothing, shoes, sticks/branches, and, on one occasion, they engaged in play with a human skull found in the cemetery (Fig. 2c). Interestingly, in 13 aggressive interactions, adult baboons shook or held sticks, branches, and stones.

**Figure 1.**
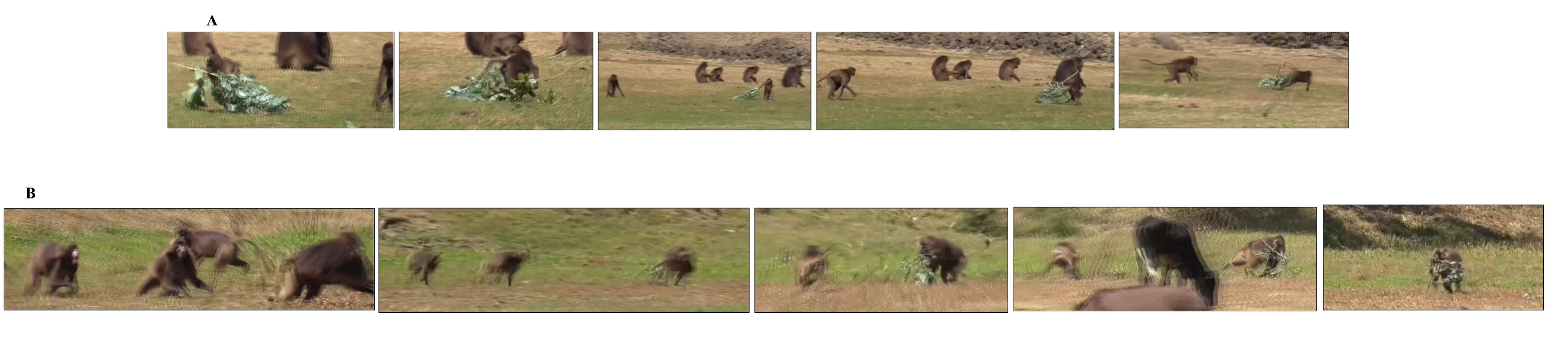
Screenshots of geladas during a social play session between immature individuals (**a**) and an aggressive interaction between adults (**b**), in both cases involving the same object present in the environment (a branch).

**Figure 2.**
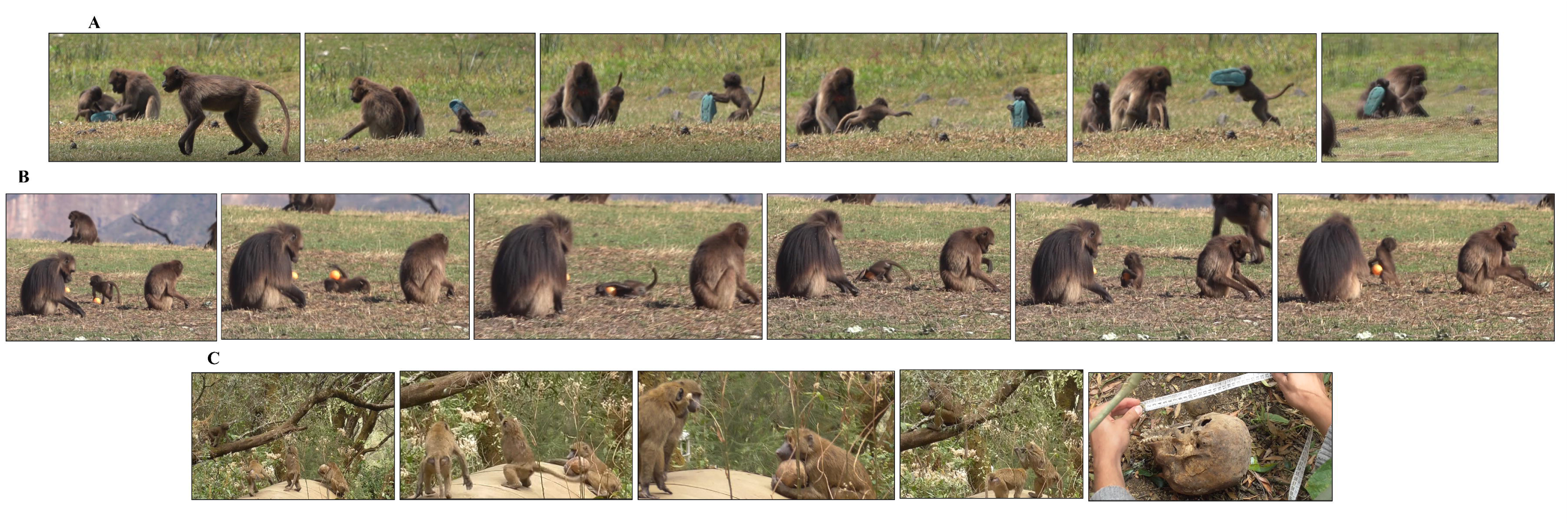
Screenshots of geladas during social object play with a blue slipper (**a**; an object naturally present in the environment) and during solitary object play with an object provided by researchers and not typically present in geladas’ environment (i.e., an orange ball) (**b**). Screenshot of olive baboons during social object play with a human skull (**c**).

The individual values of Social Object Play Index (socOPI) were higher in olive baboons than in geladas (Mann-Whitney test N_baboons_=99, N_geladas_=74, U = 2744.0, p < 0.001; Fig. 3a). There was no difference between geladas and baboons in the individual values of Solitary Object Play Index (solOPI; Mann-Whitney test N_baboons_=51, N_geladas_=20, U = 442.0, p = 0.373; Fig. 3b).

**Figure 3.**
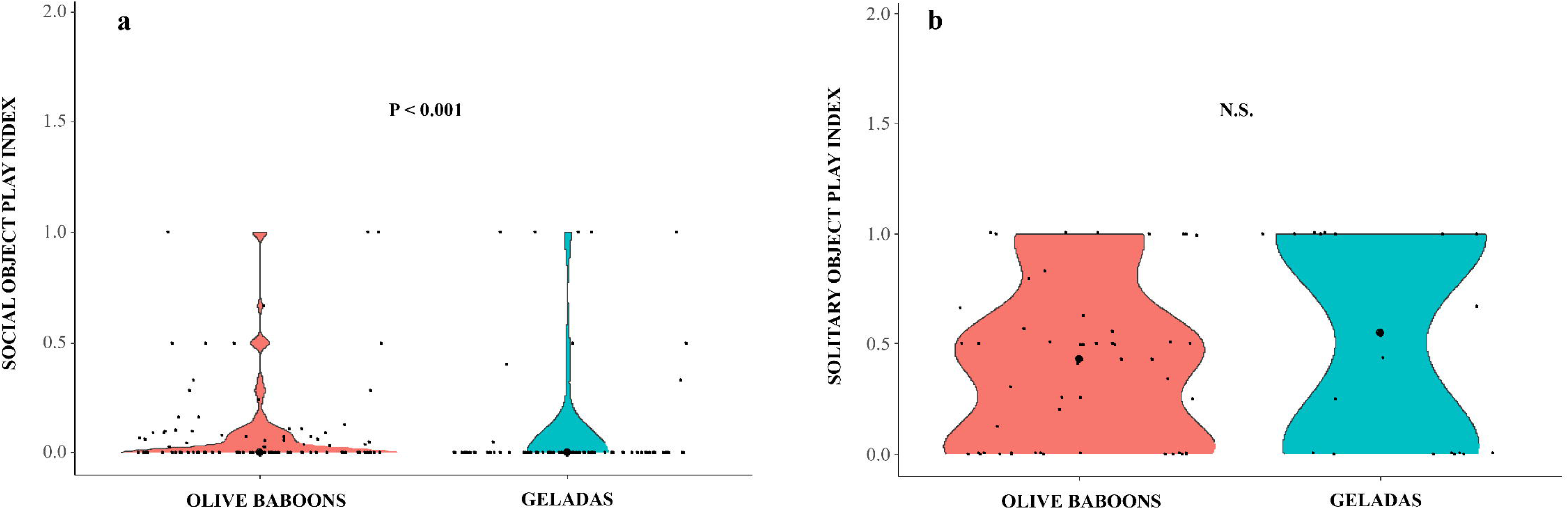
Violin plot representing the comparison between olive baboons and geladas in Social Object Play Index values (a) and Solitary Object Play Index values (b). The shape of the violin represents the density estimate of the variable considered. Dots represent the individual data points. The black central dots represent median values. Only probability of significant results is represented in the graph.

In both geladas and baboons the values of solOPI were higher than the values of socOPI (Wilcoxon test N_geladas_=13, T=1, ties=3, p = 0.006; mean value OPI ±SE: solitary 0.517 ±0.107, social 0.092 ±0.031; N_baboons_=51, T=53, ties=14, p < 0.001; mean value OPI ±SE: solitary 0.393 ±0.050, social 0.092 ±0.021). Furthermore, for geladas, also when considering only the cases where objects were provided by the researchers, the frequencies of solitary object play were higher than the frequencies of social object play (Wilcoxon test N_geladas_=34, T=145, ties=2, p = 0.014; mean value object play ±SE: solitary 1.560 ±0.514, social 0.706 ±0. 172).

## DISCUSSION

In the present study, we found that immature geladas engaged in object play – even with novel objects introduced into their environment – also in the wild and that some of the objects were used in a re-elaborated way (e.g. during aggression) by adults. The observed patterns of interactions with objects (Table 1) fall squarely within the definition of object play [1–3].

Geladas exhibited less social object play than olive baboons but, intriguingly, they showed solitary object play at levels comparable to olive baboons, a species generally described as more neophilic and object-oriented than geladas [29]. The difference in social play with objects may be interpreted in light of the distinct social systems characterizing the two species. Gelada society is characterized by high levels of tolerance within One-Male Units (OMUs) - the core social module of gelada multilevel society, composed of one adult reproductive male, several adult females, and their offspring [37–38,44,48]. For example, immature individuals engage in play at comparable levels with partners both within and outside their OMUs [49] and cohesion between OMUs is not maintained through aggressive male herding but rather through frequent affiliative interactions and tolerance among group members [37]. Conversely, baboons live in multi-male/multi-female troops marked by variable competitive relationships, conditions under which assessing relationship quality is particularly advantageous [50–53]. In olive baboons, both males and females may direct aggression toward current and potential future competitors [52,54]. These dynamics reflect a social environment in which competition and strategic relationship management are prominent features. The presence of a monopolizable object in baboons enhances the competitive component of playful interactions, as observed, for example, in playful teasing often involving repeated poking, hitting, or swinging objects in front of a partner [55]. Thus, social object play in geladas can have a less competitive function than in olive baboons when navigating social relationships [56–57].

Solitary object play represents a distinct, yet equally important, dimension of play behavior. In both human and non-human primates, object play during individual development provides opportunities to acquire knowledge about the action-relevant properties of objects, potentially facilitating the later re-elaboration of the use of such objects [57]. Through repeated interactions with objects, individuals gradually learn how these can be manipulated and employed as tools to achieve goals and solve problems [1,4,17,20,57]. In this sense, object play can be viewed as a developmental substrate for the emergence of flexible, goal-directed manipulation.

In our study, we found that geladas engaged in solitary object play at comparable levels to those observed in baboons challenging the view that baboons are the most interested in objects [29,58]. Both geladas and baboons engaged more frequently in solitary than in social object play; this trend was also observed in geladas when interacting with objects provided by researchers that are not typically present in their environment. This result is not surprising for baboons. According to the literature, baboons inhabit a wide range of environments (e.g., grasslands, savannas, mountains, rainforests) and consequently process and consume diverse types of food [33–36]. As a matter of fact, baboons have been shown to playfully interact with objects in the same way they might interact with potential food items [29]. For geladas, the result is more surprising but, on closer inspection, it can be linked to their ecological and morphoanatomical features. The persistence of arboreal behavior [39–41] is associated with the manipulation of food items other than grass, such as spherical or elliptical fruits, which we observed in the study site (e.g., prickly pears). Furthermore, geladas exhibit pronounced manual dexterity, the origins of which are often linked to the ability to solve cognitively and/or motorically demanding tasks, such as extractive foraging [59]. Skilled hand use provides a significant advantage in translating cognitive capacities into effective action. Notably, geladas maintain highly refined manual techniques even in the context of feeding [59].

Importantly, we observed immature geladas engaging in social – but mostly solitary - object play with objects available in their environment that were also used by adults during aggressive or other types of social interactions (e.g. branches; Fig. 1b; video S1).

Bergman and Kitchen [29] showed that, in the wild, geladas exhibited less neophilia and exploratory behaviors than baboons. The authors recorded the response of both species to novel objects (i.e. plastic doll, rubber ball, and metal can) and found that baboons approached, handled, and manipulated novel objects while geladas did not show either particular interest or playful interaction with these objects but simply touched or briefly picked them up, which cannot defined as object play according to its definition [1–3]. Palagi and Bergman [58] pointed out that, compared to baboons, wild geladas, are not interested in novel objects they encounter but they only glance at them and continue their foraging activity. This holds true for different study sites in the wild, but it is not extendable to all sites.

As a further step, we found that – as baboons – also wild geladas, at the Debre Libanos site, showed interest in novel objects (i.e. provided objects; Fig. 2b, video S2) that were not present in their environment. Immature geladas carried, bite, shook, threw, pulled down balls and ropes (occasionally together) or kept them in mouth or hands while rolling or pirouetting in solitary manner which falls into the object play definition (video S2; Table 1) [1–3]. Thus, these novel objects probably retain ecological relevance for this species, as both rounded shapes (e.g. fruits) and elongated shapes (e.g. vines) are found in the environment and are worth the property assessment. Also considering only the cases where the objects were provided to geladas, solitary object play remained more frequent than social object play.

In conclusion, given the relatively recent divergence between *Papio* and *Theropithecus* [30–31], together with the ecological flexibility across gelada populations, object play may reflect conserved ancestral capacities that emerge under particular circumstances. Overall, these findings caution against generalizations at the species level, as specific ecological niches and habitats can significantly influence behavior. Consequently, distinct ‘*behavioral ecotypes*’ may emerge across different populations, highlighting that the relationship between ecology, sociality, and play is more nuanced than traditionally assumed.

## Supporting information

Dataset_analysis

Video_captions_for_supporting_materials

## Ethics

The current study is non-invasive and compliant with current Ethiopian and Italian laws and University of Torino regulations, according to which no permit from the Bio-Ethical Committee was necessary. The researchers obtained the Ethiopian Wildlife Conservation Authority permission for carrying out the study.

## Data accessibility

All individual data and videos are provided as supplementary material. **Declaration of AI use**. The AI tool was not used in experimental design, data collection and analysis, and manuscript writing.

## Authors’ contributions

G.C., I.N.: conceptualization, data curation, investigation, methodology, resources, supervision, writing - review and editing; G.C.: formal analysis, writing - original draft; M.C.P., Y.Y.H., A.B., S.B.: data collection and processing; S.B.: video editing; E.D.B.: paper review, supervision; B.A.B.: supervision and project administration, paper review.

## Conflict of interest declaration

the authors declare no conflict of interests.

## Funding

this work is part of a PhD and Master thesis project supported by the Erasmus+ KA171 Partner Countries. Other fundings include: PRIN 2022 project (funded by the Italian Ministry of University and Research; MUR) “Non-verbal markers of emotional intersubjectivity: a comparative approach to empathy” (Project Code/CUP: D53C24004770006; Ministry COD. 2022ZZAZKW), CORG_RILO_24_01.

## Acknowledgments

the authors would like to thank the local guides Kabe and Hailey and the driver Brhanu for their support during field work, Uddist for restoring us with her Ethiopian coffee, the local population for assistance and collaboration.

## Notes

### Competing Interest Statement

The authors have declared no competing interest.

## REFERENCES

1. Bjorklund DF, Gardiner AK 2010 Object play and tool use: developmental and evolutionary perspectives. In The Oxford Handbook of the Development of Play (eds P Nathan, AD Pellegrini), pp. 153–171. Oxford University Press. (doi:10.1093/oxfordhb/9780195393002.013.0013)

2. Pellis SM, Burghardt GM. 2017 Play and exploration. In APA handbook of comparative psychology: Basic concepts, methods, neural substrate, and behavior (eds J Call, GM Burghardt, IM Pepperberg, CT Snowdon, T Zentall), pp. 699–722. American Psychological Association. (doi: 10.1037/0000011-034)

3. Berlyne DE 1960 Conflict, Arousal, and Curiosity. McGraw-Hill Book Company

4. Riede F, Johannsen NN, Högberg A, Nowell A, Lombard M. 2018 The role of play objects and object play in human cognitive evolution and innovation. Evol. Anthropol. 27(1), 46–59. (doi:10.1002/evan.21555)

5. Kuba MJ, Gutnick T, Burghardt GM 2014 Learning from play in octopus. In Cephalopod Cognition (eds AS Darmaillacq, L Dickel, J Mather), pp. 57–71. Cambridge University Press

6. Burghardt GM, Dinets V, Murphy JB. 2015 Highly repetitive object play in a cichlid fish (*Tropheus duboisi*). Ethology 120, 1–7. (doi: 10.1111/eth.12312)

7. Burghardt GM, Chiszar D, Murphy JB, Romano J, Walsh T, Manrod J. 2002 Behavioral complexity, behavioral development and play. In Komodo dragons: biology and conservation (eds JB Murphy, C Ciofi, C de la Panouse, T Walsh), pp. 78–117. Washington, DC: Smithsonian Press

8. Dinets V. 2015 Play behavior in crocodilians. Anim. Behav. Cog. 2(1), 49–55. (doi:10.12966/abc.02.04.2015)

9. Lambert ML, Schiestl M, Schwing R, Taylor AH, Gajdon GK, Slocombe KE, Seed AM. 2017 Function and flexibility of object exploration in kea and New Caledonian crows. R. Soc. Open Sci. 4, 170652. (doi:10.1098/rsos.170652)

10. O’Hara M, Auersperg AMI. 2017 Object play in parrots and corvids. Curr. Opin. Behav. Sci. 16, 119–125. (doi:10.1016/j.cobeha.2017.05.008)

11. Bradshaw JWS, Casey RA, Brown SL. 2012 The behaviour of the domestic cat. CABI, Boston

12. Greene WE, Melillo-Sweeting K, Dudzinski KM. 2011 Comparing object play in captive and wild dolphins. Int. J. Comp. Psychol. 24(3), 292–306. (doi:10.46867/IJCP.2011.24.03.01)

13. Torigoe T. 1985 Comparison of object manipulation among 74 species of non-human primates. Primates 26, 182–194. (doi:10.1007/BF02382017)

14. Burghardt G. 2005 The Genesis of Animal Play Testing the Limits. The MIT Press

15. Burghardt GM, Albright JD, Davis KM. 2016 Motivation, development and object play: comparative perspectives with lessons from dogs. Behaviour 153(6-7), 767–793. (doi:10.1163/1568539X-00003378)

16. Nahallage CA, Huffman MA. 2007 Age[specific functions of stone handling, a solitary[object play behavior, in Japanese macaques (*Macaca fuscata*). Am. J. Primatol. 69(3), 267–281. (doi:10.1002/ajp.20348)

17. Cenni C, Nord C, Christie JB, Wandia IN, Leca JB. 2024 How does object play shape tool use emergence? Integrating observations and field experiments in longtailed macaques. Anim. Behav. 218, 239–254. (doi: 10.1016/j.anbehav.2024.09.001)

18. Bapat A, Kempf AE, Friry S et al. 2025 Patterns of object play behaviour and its functional implications in free-flying common ravens. Sci. Rep. 15, 137. (doi:10.1038/s41598-024-83856-9)

19. Reader SM, Laland KN 2002 Social intelligence, innovation, and enhanced brain size in primates. Proc. Nal. Acad. Sci. 99 (7), 4436–4441. (doi:10.1073/pnas.062041299)

20. Riede F, Walsh MJ, Nowell A, Langley MC, Johannsen NN. 2021 Children and innovation: Play, play objects and object play in cultural evolution. Evol. Hum. Sci. 3, e11. (doi:10.1017/ehs.2021.7)

21. Bădescu I, McKerracher LJ, Sellen DW, et al. 2025 Atypical tool and object use in wild immature chimpanzees reveals developmental pathways to innovation. Sci. Rep. 15, 36396. (doi:10.1038/s41598-025-20487-8)

22. Ramsey JK, McGrew WC. 2005 Object play in great apes. Studies in Nature and Captivity. In The nature of play: Great apes and humans (eds AD Pellegrini, PK Smith), pp. 89-112. The Guilford Press

23. Leca JB, Gunst N, Huffman MA. 2007 Japanese macaque cultures: Interand intra-troop behavioural variability of stone handling patterns across 10 troops. Behaviour 144, 251–281.

24. Jordan EJ, Völter CJ, Seed AM. 2022 Do capuchin monkeys (*Sapajus apella*) use exploration to form intuitions about physical properties? Cog. Neuropsychol. 38(7-8), 531e543. (doi:10.1080/02643294.2022.2088273)

25. Huffman MA, Quiatt D. 1986. Stone handling by Japanese macaques (*Macaca fuscata*): Implications for tool use of stone. Primates 27, 413e423. (doi:10.1007/bf02381887)

26. Huffman MA. 1996 Acquisition of innovative cultural behaviors in non-human primates: a case study of stone handling, a socially transmitted behavior in Japanese macaques. In Social learning in animals: The roots of culture (eds CM Heyes, BG Galef Jr.), pp. 267–289. Academic Press. (doi:10.1016/b978-012273965-1/50014-5)

27. Leca JB, Gunst N. 2023 The exaptive potential of (object) play behavior. Int. J. Play 12, 40–52. (doi:10.1080/21594937.2022.2152184)

28. Westergaard GC. 1992 Object manipulation and the use of tools by infant baboons (*Papio cynocephalus anubis*). J. Comp. Psychol. 106(4), 398–403. (doi:10.1037/0735-7036.106.4.398)

29. Bergman TJ, Kitchen DM. 2009 Comparing responses to novel objects in wild baboons (*Papio ursinus*) and geladas (*Theropithecus gelada*). Anim. Cogn. 12, 63–73. (doi:10.1007/s10071-008-0171-2)

30. Jablonski NG. 2005 Theropithecus: the rise and fall of a primate genus. Cambridge: Cambridge University press

31. Gilbert CC, Frost SR, Pugh KD, Anderson M, Delson E. 2018 Evolution of the modern baboon (*Papio hamadryas*): a reassessment of the African Plio-Pleistocene record. J. Hum. Evol. 122, 38–69. (doi:10.1016/j.jhevol.2018.04.012)

32. Cangiano M, Palagi E. 2020 First evidence of stone handling in geladas: from simple to more complex forms of object play. Behav. Proc. 180, 104253. (doi:10.1016/j.beproc.2020.104253)

33. Altmann SA. 1998 Foraging for survival: Yearling baboons in Africa. Chicago: University of Chicago Press

34. Okecha AA, Newton-Fisher NE. 2006 The diet of olive baboons (*Papio anubis*) in the Budongo Forest Reserve, Uganda. In Primates of western Uganda. Developments in Primatology: Progress and Prospects (eds NE Newton-Fisher, H Notman, JD Paterson, V Reynolds), pp. 61–73. New York, NY: Springer New York

35. Barrett L, Henzi SP. 2008 Baboons. Curr. Biol. 18(10), 404–406. (doi: 10.1016/j.cub.2008.02.074)

36. Kifle Z. 2021 Human-olive baboon (*Papio anubis*) conflict in the human-modified landscape, Wollo, Ethiopia. Glob. Ecol. Conserv. 31, e01820. (doi:10.1016/j.gecco.2021.e01820)

37. Dunbar RIM, Dunbar P. 1975 Social Dynamics of Gelada Baboons. Karger, Basel

38. Snyder-Mackler N, Beehner JC, Bergman TJ. 2012 Defining higher levels in the multilevel societies of geladas (*Theropithecus gelada*). Int. J. Primatol. 33(5), 1054–1068. (doi:10.1007/s10764-012-9584-5)

39. Kifle Z, Bekele A. 2021 Feeding ecology and diet of the southern geladas (*Theropithecus gelada obscurus*) in human-modified landscape, Wollo, Ethiopia. Ecol. Evol. 11, 11373–11386. (doi:10.1002/ece3.7927)

40. Kifle Z. 2023 The mystery of gelada (*Theropithecus gelada*) survival and adaptation in the highly anthropogenically-modified landscapes in the Ethiopian highlands: a review. Glob. Ecol. Conserv. 47, e02669. (doi:10.1016/j.gecco.2023.e02669)

41. Fleagle JG, Baden AL, Gilbert CC. 2025 Primate Adaptation and Evolution. Academic Press

42. Maier W. 1993 Adaptations in the hands of cercopithecoids and callitrichids. In Hands of Primates (eds H Preuschoft, DJ Chivers), pp. 191–198. Vienna: Springer. (doi:10.1007/978-3-7091-6914-8_12)

43. Fashing PJ, Nguyen N, Venkataraman VV, Kerby JT. 2014 Gelada feeding ecology in an intact ecosystem at Guassa, Ethiopia: variability over time and implications for theropit and hominin dietary evolution. Am. J. Phys. Anthropol. 155, 1–16. (doi:10.1002/ajpa.22559)

44. Caselli M, Zanoli A, Dagradi C, Gallo A, Yazezew D, Tadesse A, Capasso M, Ianniello D, Rinaldi L, Palagi E, Norscia I. 2021 Wild geladas (*Theropithecus gelada*) in crops-more than in pasture areas-reduce aggression and affiliation. Primates 62(4), 571–584. (doi:10.1007/s10329-021-00916-8)

45. Pedruzzi L, Francesconi M, Galotti A, Bogale BA, Palagi E, Lemasson A. 2025 Wild gelada monkeys detect emotional and prosocial cues in vocal exchanges during aggression. PLoS One 20(5), e0323295. (doi:10.1371/journal.pone.0323295)

46. Altmann J. 1974 Observational study of behavior: sampling methods. Behaviour 49(3-4), 227–266. (doi:10.1163/156853974x00534)

47. Cordoni G, Ciarcelluti G, Pasqualotto A, Perri A, Bissiato V, Norscia I. 2023 Is it for real? Structural differences between play and real fighting in adult chimpanzees (*Pan troglodytes*). Am. J. Primatol. 85(9), e23537. (doi:10.1002/ajp.23537)

48. Dunbar RIM. 1992 A model of the gelada socio-ecological system. Primates 33, 69–83. (doi:10.1007/BF02382763)

49. Gallo A, Caselli M, Norscia I, Palagi E. 2021 Let’s unite in play! Play modality and group membership in wild geladas. Behav. Proc. 184, 104338. (doi:10.1016/j.beproc.2021.104338)

50. Silk JB, Beehner JC, Bergman TJ, Crockford C, Engh AL, Moscovice LR, Wittig RM, Seyfarth RM, Cheney DL. 2009 The benefits of social capital: close social bonds among female baboons enhance offspring survival. Proc. R. Soc. B 276,3099–3104. (doi:10.1098/rspb.2009.0681)

51. Silk JB, Beehner JC, Bergman TJ, Crockford C, Engh AL, Moscovice LR, Wittig RM, Seyfarth RM, Cheney DL. 2010 Strong and consistent social bonds enhance the longevity of female baboons. Curr. Biol. 20, 1359–1361. (doi:10.1016/j.cub.2010.05.067)

52. Silk JB, Roberts ER, Barrett BJ, Patterson SK, Strum SC. 2017 Female–male relationships influence the form of female–female relationships in olive baboons, *Papio anubis*. Anim. Behav. 131, 89–98. (doi:10.1016/j.anbehav.2017.07.015)

53. Patterson SK, Strum SC, Silk JB. 2022 Early life adversity has long-term effects on sociality and interaction style in female baboons. Proc. R. Soc. B 289, 20212244. (doi:10.1098/rspb.2021.2244)

54. MacCormick HA, MacNulty DR, Bosacker AL, Lehman C, Bailey A, Collins DA, Packer C. 2012 Male and female aggression: lessons from sex, rank, age, and injury in olive baboons. Behav. Ecol. 23(3), 684–691. (doi:10.1093/beheco/ars021)

55. Winkler SL, Cartmill EA. 2026 Does playful teasing help great apes learn about social relationships? Philos. Trans. R Soc. Lond. B Biol. Sci. 381(1943), 20240371. (doi:10.1098/rstb.2024.0371)

56. Cordoni G, Norscia I. 2024 Nuancing ‘emotional’ social play: does play behaviour always underlie a positive emotional state? Animals 14(19), 2769. (doi:10.3390/ani14192769)

57. Parker ST, Gibson KR. 1977 Object manipulation, tool use and sensorimotor intelligence as feeding adaptations in Cebus monkeys and great apes. J. Hum. Evol. 6(7), 623–641. (doi:10.1016/S0047-2484(77)80135-8)

58. Palagi E, Bergman TJ. 2021 Bridging captive and wild studies: behavioral plasticity and social complexity in *Theropithecus gelada*. Animals 11(10), 3003. (doi:10.3390/ani11103003)

59. Truppa V, Gamba M, Togliatto R, Caselli M, Zanoli A, Palagi E, Norscia I. 2024 Manual preference, performance, and dexterity for bimanual grass[feeding behavior in wild geladas (*Theropithecus gelada*). Am. J. Primatol. 86(5), e23602. (doi:10.1002/ajp.23602)

