## Supplementary material for "FIRST EVIDENCE OF OBJECT PLAY IN WILD GELADAS: FUNCTIONAL IMPLICATIONS FOR LATER UTILITY AND RE-ELABORATED OBJECT USE IN ADULTHOOD": Video_captions_for_supporting_materials

**Video S1 – Interactions with objects: play *vs* aggression**

Video clips describing (i) a social object-play interaction with a branch between two immature individuals, and (ii) an aggressive interaction between an adult male and adult females, in which the male uses a branch. Video editing Sebastiano Bergamo.

**Video S2 – Solitary and social object play**

Video clips describing (i) an infant engaging in a solitary object-play session involving an object (an orange ball) provided by the researchers and not naturally present in the geladas’ environment, and (ii) a social object-play session involving a slipper between two infants.
